# A Versatile Robotic Platform for the Design of Natural, Three-Dimensional Reaching and Grasping Tasks in Monkeys

**DOI:** 10.1101/653840

**Authors:** B Barra, M Badi, MG Perich, S Conti, SS Mirrazavi Salehian, F Moreillon, S Wurth, M Kaeser, P Passeraub, T Milekovic, A Billard, S Micera, M Capogrosso

## Abstract

Translational studies on motor control and neurological disorders require detailed monitoring of sensorimotor components of natural limb movements in relevant animal models. However, available experimental tools do not provide a sufficiently rich repertoire of behavioral signals. Here, we developed a robotic platform that enables the monitoring of kinematics, interaction forces, and neurophysiological signals during user-definable upper limb tasks for monkeys. We configured the platform to position instrumented objects in a three-dimensional workspace and provide an interactive dynamic force-field. We show the relevance of our platform for fundamental and translational studies with three example applications. First, we study the kinematics of natural grasp in response to variable interaction forces. We then show simultaneous and independent encoding of kinematic and forces in single unit intra-cortical recordings from sensorimotor cortical areas. Lastly, we demonstrate the relevance of our platform to develop clinically relevant brain computer interfaces in a kinematically unconstrained motor task

## INTRODUCTION

Modern approaches to system neuroscience as well as the study of motor recovery in clinical applications require detailed characterization of sensorimotor neural activity underlying natural, three-dimensional movements rather than artificially constrained behaviours. In this context, an ideal experimental set-up should provide 1) flexible and instrumented workspaces to allow natural, but reproducible motor behaviours and 2) multimodal electrophysiology recordings and behavioural signals integration.

However, to date, no experimental set-up allows the integration of electrophysiological recordings with extrinsic signals quantifying natural interactions with the environment.

Indeed, several groups developed sophisticated experimental platforms capable of characterizing unconstrained kinematics but did not provide information on interaction forces (Churchland et al., 2012; Hu et al., 2018; Schaffelhofer and Scherberger, 2016; Schwartz et al., 1988; Umeda et al., 2019; Vargas-Irwin et al., 2010). Conversely, the study of active force control historically focused on constrained tasks in restricted experimental settings, often employing only 1-degree of freedom movements (Cheney and Fetz, 1980; Ethier et al., 2012; Herter et al., 2009; Moritz et al., 2007; Nishimura et al., 2013; Seki et al., 2003). These limitations arise from the difficulty of pairing unconstrained natural movements (Schwartz et al., 1988) with instrumented workspaces that allow the execution of active motor tasks (Cheney and Fetz, 1980) while measuring force interactions with the surrounding environment.

We believe that functionalized workspaces that promote natural movements could be designed by extending the concept of classical planar robotic tasks (de Haan et al., 2018; London and Miller, 2013; Omrani et al., 2016) to three-dimensional workspaces. Such setup could provide detailed measurement of kinematics and interaction forces throughout actions involving reaching, grasping and manipulation of objects. At the same time, it would enable complete freedom in defining the spatial constraints, force fields and perturbations of tasks that resembles natural, three-dimensional movements.

Here, we present a versatile robotic platform that combines neurophysiological, mechanical, and kinematic measurements within a customizable three-dimensional experimental environment. This platform consists of 1) a seven degrees-of-freedom compliant robotic arm (LBR iiwa, KUKA, Augsburg, Germany) 2) a custom software control package 3) force and grip pressure sensors integrated in the robot and 4) modules for data synchronisation.

To demonstrate the potential of our platform for both basic and translational studies in motor control, we programmed the robotic arm to present instrumented objects to trained monkeys in a three-dimensional workspace and oppose elastic resistance to displacements of the end effector. By pairing this system with intra-cortical neural recordings of sensorimotor areas, we created an instrumented platform that provides a rich portfolio of signals for the investigation of natural motor behaviour.

We trained two monkeys to reach for the robot and pull the end effector to receive a food reward. We first demonstrate the performance and safety of our platform. We then use the platform to study the kinematic and dynamic components of movement and how these vary when applying different strength of dynamic elastic resistance to the target object movement. Third, we show that activity of neurons in both the motor and somatosensory areas encode specific components of the task such as force, kinematics or object contact. Fourth, we argue that our framework can be instrumental to neural engineering studies seeking to decode movement information from motor and sensory areas during natural behaviour.

Our versatile control structure allows for the design and implementation of a variety of tasks that can support both fundamental and translational studies of motor control.

## RESULTS

### A versatile robotic framework

Our platform (Figure 1) consists of 1) a seven-degree-of-freedom robotic arm (Intelligent Industrial Work Assistant, IIWA - KUKA, Augsburg, Germany), 2) a custom built software package that enables closed-loop control of the robot arm, 3) a synchronized interaction force recording system, 4) a strain-gauge grip pressure sensor, 5) an infrared video tracking system to measure three-dimensional joint kinematics (Vicon, Oxford, UK) and an 6) electrophysiology system (Blackrock Microsystems, Salt Lake City, USA). We assessed the versatility and efficacy of our framework by programming a robotic task for monkeys. We configured the robot to position objects in a three-dimensional workspace and trained monkeys to reach and pull on the objects while kinetic, kinematic and neural signals were simultaneously recorded.

**Figure 1.**
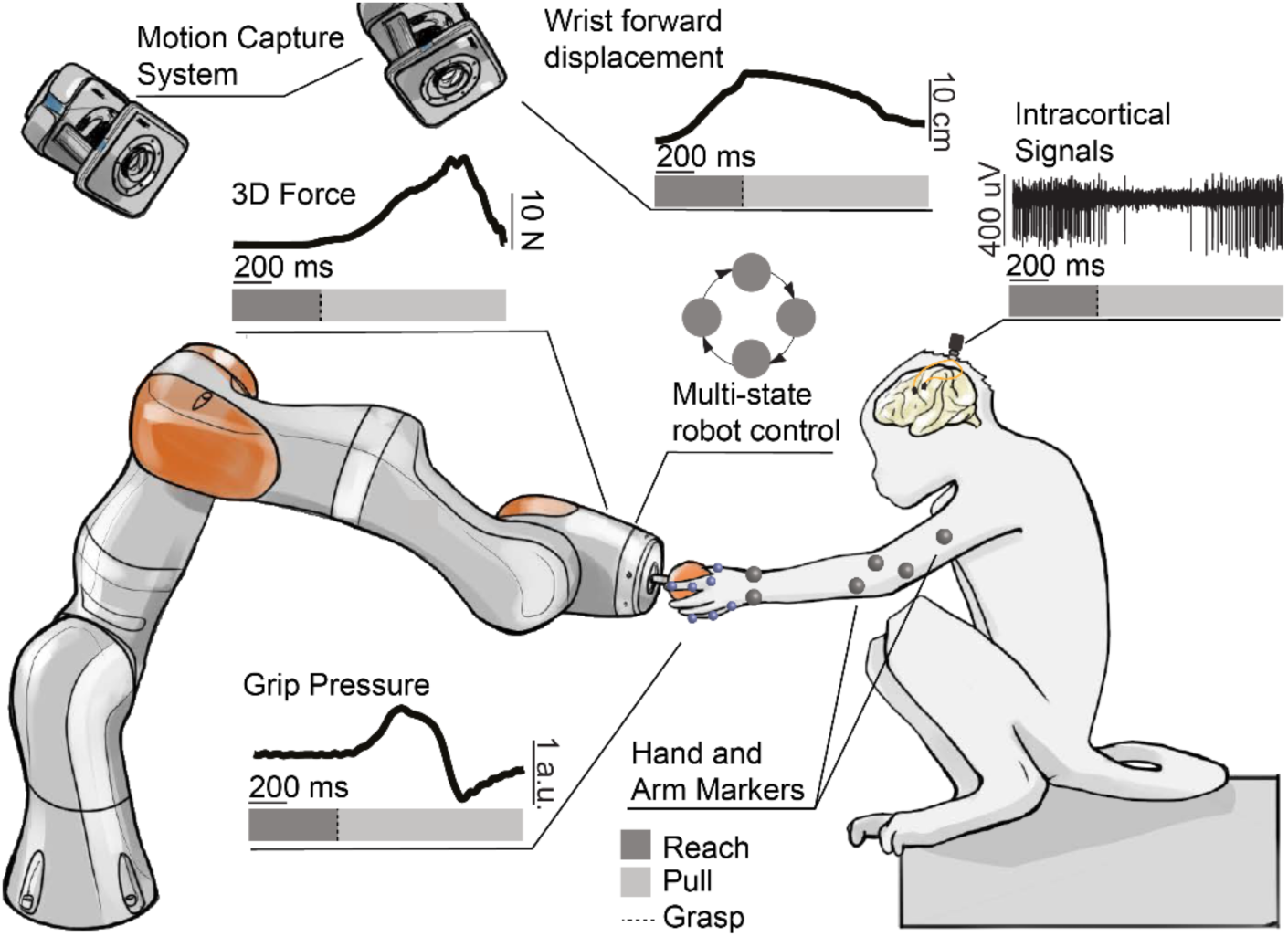
Robotic framework for the study of reaching and grasping. A monkey is implanted with microelectrode arrays in the arm and hand sensorimotor areas. A robotic arm is used to present target objects to the monkey in different configurations of the 3D space. The animal can interact with the robotic arm by reaching for, grasping and pulling on the target objects. A multi-state user-definable robot control software is implemented to allow simple task re-configuration. The 3D force applied by the monkey on the robotic joints is measured in real time. The target objects are instrumented with a pressure sensor able to measure the applied grip pressure. A motion capture system is used to track the monkey arm and finger 3D positions using reflective markers. Multiunit brain activity is recorded through a high-resolution electrophysiology system.

#### Closed-Loop control infrastructure

The IIWA robotic arm features a large workspace (Figure S1) allowing ranges of motion that are compatible with both human and monkey reaches. Additionally, the robotic arm is able to actively lift up to 7Kg of weight, which makes it robust to manipulation by monkeys.

We developed a software package that implements a real-time closed-loop control (Figure 2A, <u>10.5281/zenodo.3234138</u>) configured as a finite state machine. This allows fast configuration of tasks that proceed through several phases, where each phase requires a different behaviour of the robot.

**Figure 2.**
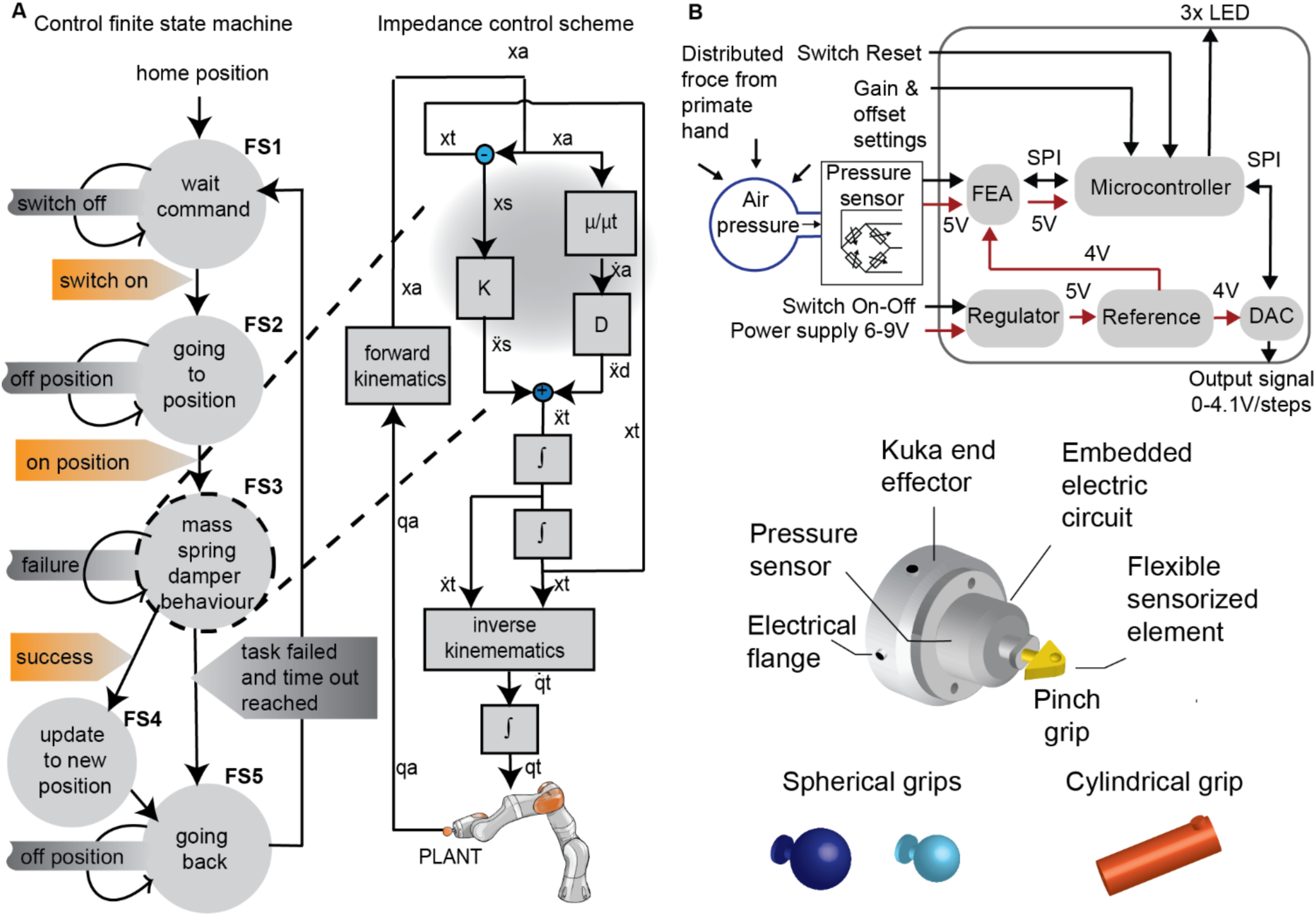
Robotic arm control scheme and grip pressure sensor design. **(A)** Finite State Machine of the robotic control during the behavioural task. When a trial starts, the robot moves the end effector to a pre-determined position in space using standard impedance joint control strategy. When the position is reached, the finite state machine switches to a custom implemented impedance control mode to allow interaction with the animal (right panel). In this modality, the end effector behaves like a mass-spring-damper system whose parameters are entirely definable by the user and can be easily modified. xa, ẋa, = measured position and velocity in cartesian space; xs, ẍs = spring derived position and acceleration in cartesian space; ẍd = damping derived acceleration in cartesian space; xt, ẋt, ẍt = target position, velocity and acceleration in cartesian space; qt, qt = target position and velocity in the joint space. **(B)** Top: pressure sensor electronic circuit design describing the electronic components, the communication protocols and the voltage input values. Bottom: schematic of the pressure sensor assembly fixed at the end effector of the robot with 3D drawings of the spherical, cylindrical and pinch-like objects. DAC: digital-to-analog converter - FEA: front-end amplifier – LED: light-emitting diode - SPI: serial peripheral interface.

In our specific example application, at the beginning of the trial, the robot moves the end effector to a pre-determined position in space using impedance joint control. Upon reaching position, the state machine switches to mass-spring damper behaviour (Figure 2A). Stiffness and damping parameters are definable by the user. Variations from this behaviour can be easily configured using our software.

#### Omnidirectional measurement of grip strenght

We designed an air-pressure-based gauge sensor that can measure the applied grip force independently from an object shape in the interval −0.7 kPa to 40 kPa. We 3-D-printed hollow objects of different shape and size (Figure 2B) and connected to a steel enclosure hosting the sensor. In this configuration the grip strength is proportional to the air flux produced by compression of the object. We designed a programmable electronic circuit (Figure 2B) that digitizes pressure measurements while power and data transfer are provided through connection ports located at the robot end effector thereby limiting external wires at the subject-robot interface.

#### System performance

In order to assess the safety and usability of the robotic platform during animal interaction, we first ran a series of tests on a naïve human subject. The subject was instructed to wait for a sound cue, and then reach for, grasp and pull an object mounted on the end effector. Our close-loop control scheme allowed the robot to comply with the movements imposed by the subject while opposing an elastic force. The three-dimensional pulling force was continuously monitored and recorded. When the object was pulled across a predetermined virtual border, the robot returned to a starting position where it waited for the next command (Figure 2A, **Video 1**).

To validate the reproducibility of this behaviour we computed the *positioning error* as measured by the distance between the target position and the actual position reached by the robot (Figure 3A). We also tested the ability of the robot to consistently hold a specified position over time (*drop error,* Figure 3A). All measures have been repeated on n = 14 positions in the workspace. *Positioning errors* were on average below 1.1% of the reachable workspace (0.186±2.4 mm, −0.133±6.5 mm and 0.008±8.5 mm for the x, y, and z direction, respectively, Figure 3B). Our analysis revealed no relationship between positioning error and end effector spatial location (Figure 3C). Similarly drop errors were uniform across space and below 0.4 % of the reachable workspace (1.0±1.8 mm, 0.8±2.2 mm and 2.1±0.3 mm for the x, y, and z direction). Measured forces were consistent across sessions (n=2 sessions) (Figure 3E).

**Figure 3.**
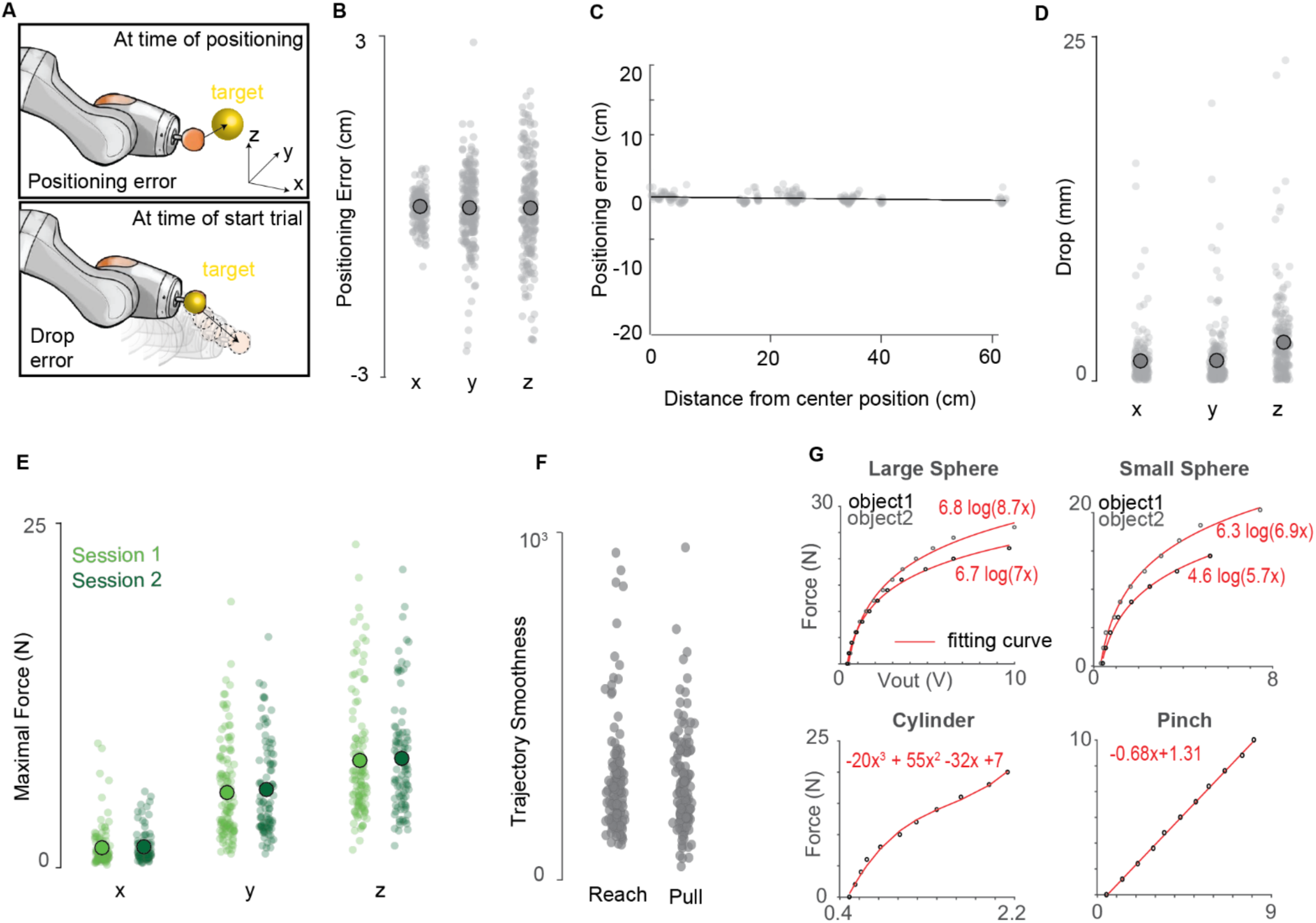
Evaluation of the system performances: **(A)** Graphical representation of the measures of reproducibility of the three-dimensional positioning. The positioning error is the distance between the set and measured robot end effector position; the drop error is the distance between the measured robot end effector position at the cue-time and the measured robot endpoint position 0.4 seconds after the cue-time. **(B)** Positioning error on the x, y and z dimension **(C)** Positioning error values as a function of the distance from the center of the robot workspace. Linear regression *α*= −0.009, RMSE = 0.05. **(D)** Drop error on the x, y and z dimension. **(E)** Maximal force exerted on the robot end effector on the x, y and z axis. **(F)** Comparison of maximal force values on the x, y and z axis for 2 different sessions. **(G)** Comparison of trajectory smoothness values during the reach (robot not interacting with the subject) and pull (robot is interacting with the subject). Data presented here were collected over 2 sessions of n=100 trials each. **(H)** Calibration curves describing the voltage to pressure relationship for each object geometry (n=1 measure per point).

We next evaluated the robot compliance upon interaction with a monkey. In particular, we verified that passive movements of the end effector did not introduce sudden and unexpected perturbations of the arm kinematic trajectories when pulling on the object.

We then trained Mk-Cs to perform the same task described above while we measured arm joint kinematics (Figure 1). We computed the smoothness index (Hans-Leo Teulings et al., 1997) of arm joints and end effector trajectories during both the reach and object interaction (pull) phases. The smoothness during the pulling phase was comparable to that computed in reaching phase (Figure 3F) suggesting that the robot did not affect the dynamics of natural arm movements.

Finally, we characterized the force to voltage relationship changes in grip pressure sensor for different object geometries. Calibration curves showed that spherical and cylindrical objects exhibit a non-linear voltage/pressure behaviour while the pinch-like geometry displayed a linear correlation between applied force and measured voltage (Figure 3G).

### Applications to basic and translational studies

We investigated the kinematic, kinetic and neural components of natural three-dimensional reaching, grasping and pulling behaviour in monkeys. Two adult *Macaca fascicularis* female monkeys (Mk-Jo and Mk-Cs) were implanted with a pair of 64-channel microelectrode arrays (Blackrock Microsystems, Salt Lake City, UT, USA). The arrays were placed in the hand and arm area of the right primary motor (M1) and right primary somatosensory (S1, area 1/2) cortices (London and Miller, 2013; Pons et al., 1985) (Figure 4A). Brain signals were synchronized to interaction forces, grip pressure and arm and hand kinematics. We implemented two tasks. In both the animal reached freely towards an object, performed a specific grasp, and compensated for the dynamic resistance applied by the robot to pull the object across a virtual border. In Task 1 the monkey reached for four objects of different shapes: a small sphere, a large sphere, a cylinder, and a “pinch” object. Each object encouraged the animal to use a specific grasp: three finger grip (small sphere), whole hand grip (large sphere), two-finger precision grip (pinch), and power grip (cylinder) (Figures 4B and 5A). The objects were presented at the same position. The joint impedance varied across “low”, “medium” and “high” level of resistance to object displacement. In Task 2, the monkey reached for the small sphere presented at different positions in space (“central”, “left”, or “right”, Figure 4C).

**Figure 4.**
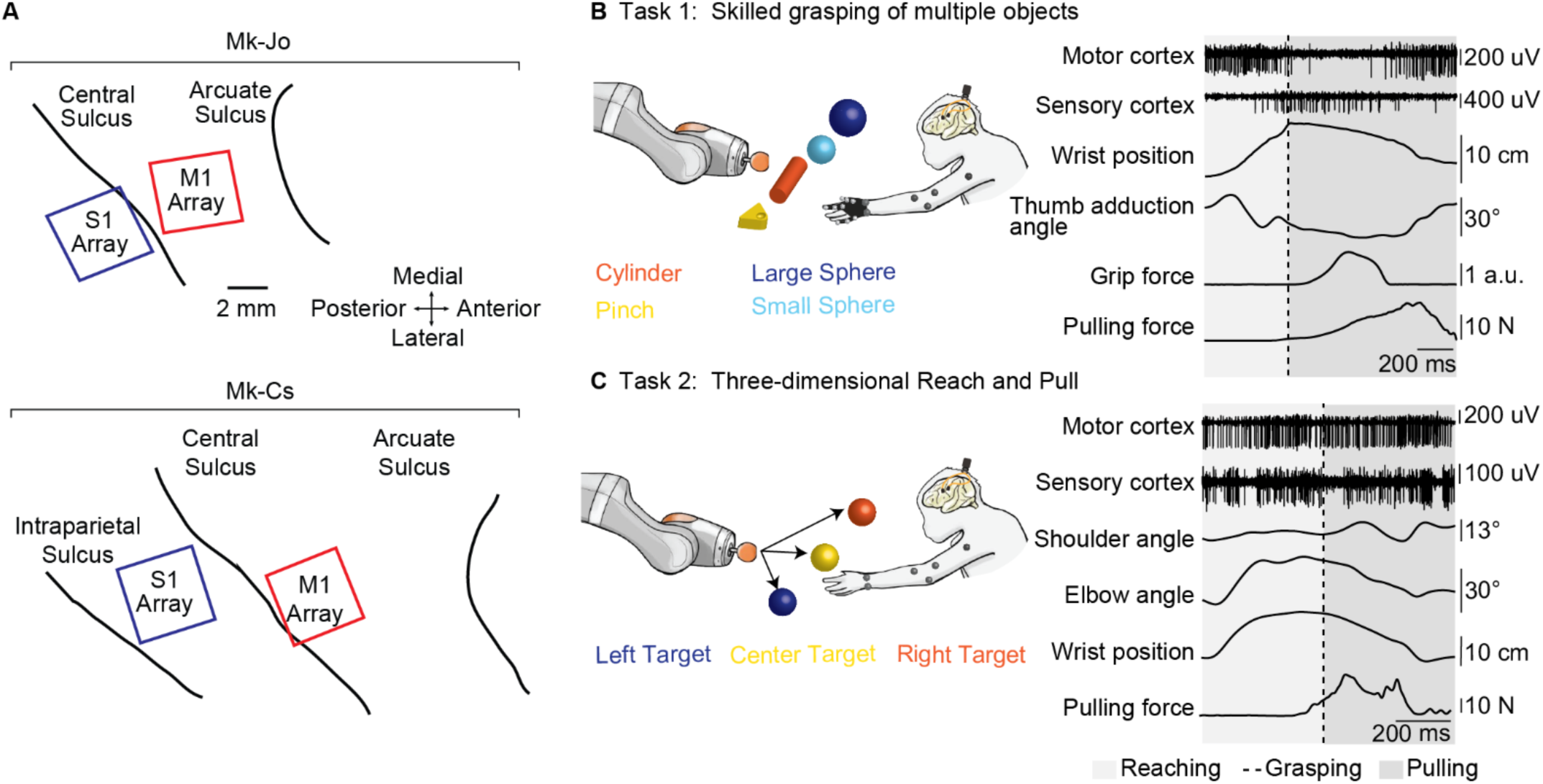
Multimodal electrophysiology during unconstrained reaching and grasping. **(A)** Utah arrays placement for Mk-Jo and Mk-Cs. Each animal received two arrays of 64 channels. M1 hand and arm areas where identified in Mk-Jo and Mk-Cs respectively through intra-operative electrical stimulation. **(B)** Schematic of a skilled grasping task during which the monkey had to reach for different types of objects and pull them towards a pre-determined spatial threshold. **(C)** Schematic of the 3D reaching task during which the monkey had to reach for an object placed at different positions in space and pull it towards a pre-determined spatial threshold. Examples of synchronous multiunit neural recordings, hand and arm kinematics, grip pressure and pulling force recordings are shown on the right for both tasks.

**Figure 5.**
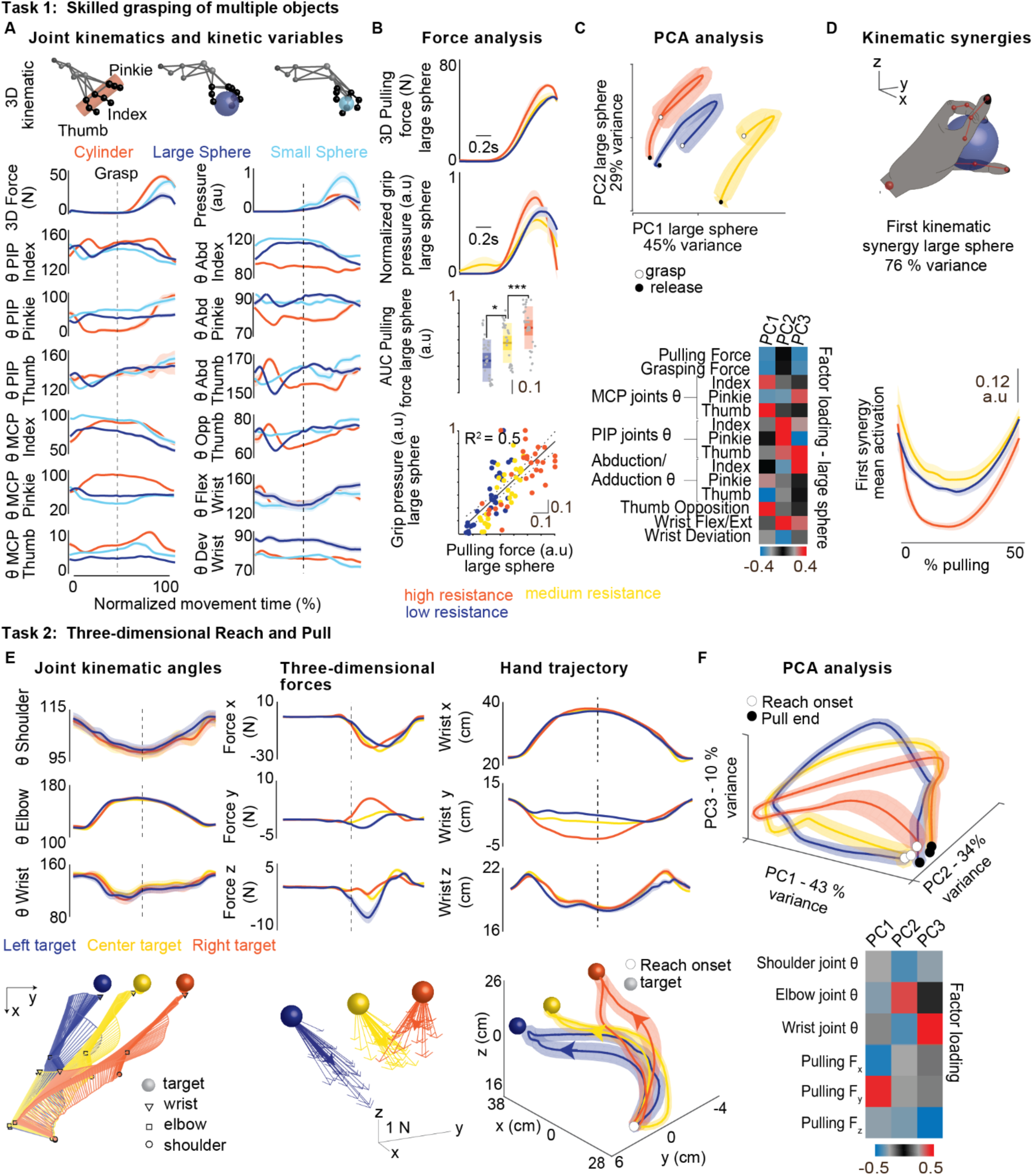
Arm and hand dynamics during natural reaching and grasping. **(A)** Examples of kinematic and dynamic variables for three different objects (n_=_15 trials per object, one recording session). 3D hand configuration is shown on top for each object. **(B)** Example of normalized pulling force and grip pressure obtained for three levels of resistance when pulling on the large spherical object (n=10 trials per resistance condition, one recording session). The box-plots the median, the 1.96 SEM and the STD of the pulling force area under the curve (AUC) during the movement (n=30 trials per condition, three recording sessions). Data were normalized by the maximal pulling force measured within each session. Increase in resistance had a significant effect on the pulling force impulse (Kruskal-Wallis, p <0.001 with post hoc correction, * p_low-med_ <0.05, *** p_low-high_ <0.001, *** p_med-high_ <0.001). Correlation analysis between the grip pressure and the pulling force impulses (AUC) revealed a linear correlation between these two variables (R^2^=0.5 and p_value_ = 1.6×10^-16^, n=30 trials per condition, three recording sessions). **(C)** Principal component analysis of the kinematic and dynamic components of the movement for the three levels of mechanical resistances (n=10 trials per condition, large sphere object, one recording session). The color-coded representation of factor loadings identifies variables that contributed most to the differences observed between the difference levels of resistance. **(D)** Top panel: postural kinematic synergy defined by the first principal component. Bottom panel: Time course activation of the first PC averaged over trials shows modulation for high level of resistance compared to low and medium resistances, indicating a change in kinematic grasping strategy to overcome increased levels of resistance (n=10 trials per condition, large sphere object, one recording session). **(E)** Examples of kinematic and dynamic variables for 3D reaching towards different spatial targets (n_=_25 trials per position, one recording session). Joint kinematics angles, three dimensional pulling force and wrist trajectories are shown for the left, center and right position. Stick diagram depicts evolution of the arm joint angles for the three conditions while maxima pulling force is represented as 3D vectors. Thick stroke arrows represents the average maximal force value. Wrist trajectories are also plotted in 3D. **(F)** Principal component analysis of the kinematic and kinetic components of the movement reveals partial overlapping of the movement performed to reach the three different target positions (n_=_25 trials per position, one recording session). Color-coded representation of factor loadings identifies the x and y components of the pulling force as the most meaningful features for the first PC while the wrist and elbow joints associate with high loadings in the second and third PC respectively. Thicker lines represents the data mean while shaded areas depicts the SEM.

#### Arm and hand dynamics during natural reaching and grasping

We first explored how kinematic and kinetic components of movement varied during natural reach and grasp paradigms, and how these evolved under different levels of resistance. The animal adapted the hand configuration according to the selected objects (Figure 5A). Fingers joint angles showed closer motor patterns for the small and large sphere than for the cylindrical object. Interestingly, grip pressure was larger for the small spherical grip than for the other objects, presumably because strong grip was required to compensate for the small surface of hand contact. Grasping pressure and pulling forces increased in response to an increase in resistance (Figure 5B, **Figure S3B**), indeed during pulling grip pressure was linearly correlated to pulling force (Figure 5B, R^2^ = 0.5). We performed a principal component analysis (PCA) of kinetic and kinematic features of grasping. This analysis revealed that grasping features evolved over well clustered smooth trajectories for different resistance levels during movement (Figure 5C, Figure S2C).

The two leading principal components showed strong correlations to finger joint kinematic features, as well as kinetic features (Figure 5C). This suggested that the monkey adapted its kinematic strategy to overcome higher resistances. To verify this hypothesis, we extracted kinematic synergies (Mason et al., 2001; Santello et al., 1998) (Figure 5D, Figure S2D). We found that the activation of the main synergy (first component, 70% of variance for all objets Figure 5D, Figure S2D) changed across resistance levels for spherical objects, but not for the cylinder (Figure 5D, Figure S2D). This suggests that the monkey adapted the grasping strategy to execute higher force levels. In the case of the cylindrical grip, the monkey overcame large resistance levels by generating stronger puling torque without substantial change in his grip pattern.

We then inspected kinematic behaviour for different spatial targets in Task 2 (Figure 5E). Joint angles showed a very consistent motor pattern across the different spatial positions, even if the animal was not trained to follow a specific strategy and was let free to reach the object without time or spatial constraints. Forces along the pulling direction were similar across all the target positions, confirming that the effort needed to pull the robot end effector past the threshold on the x-axis was proportional to the end effector displacement from the target position. Forces along the y-axis were markedly different, suggesting that the monkey always displaced the end effector towards its body center when pulling.

Projections in the PC space formed smooth and separated trajectories that spanned the three-dimensional manifold defined by first PCs (Figure 5F). Most of the variance was explained by the force exerted in the y direction as suggested by the pulling force profiles (Figure 5E).

#### Sensorimotor neural dynamics during natural reaching and pulling

We then used our robotic framework to study the neural activity in M1 and S1 during natural reaching and grasping. Nearly all channels showed high modulation of multi-unit firing rates for both sensory and motor areas in both monkeys (Figure 6A and C). Interestingly, activity arising from the arm somatosensory area (Mk-Cs) was strongly modulated during the whole movement while the largest response in the somatosensory area of the hand (Mk-Jo) occurred shortly following the grasp (Figure 6A and C).

**Figure 6.**
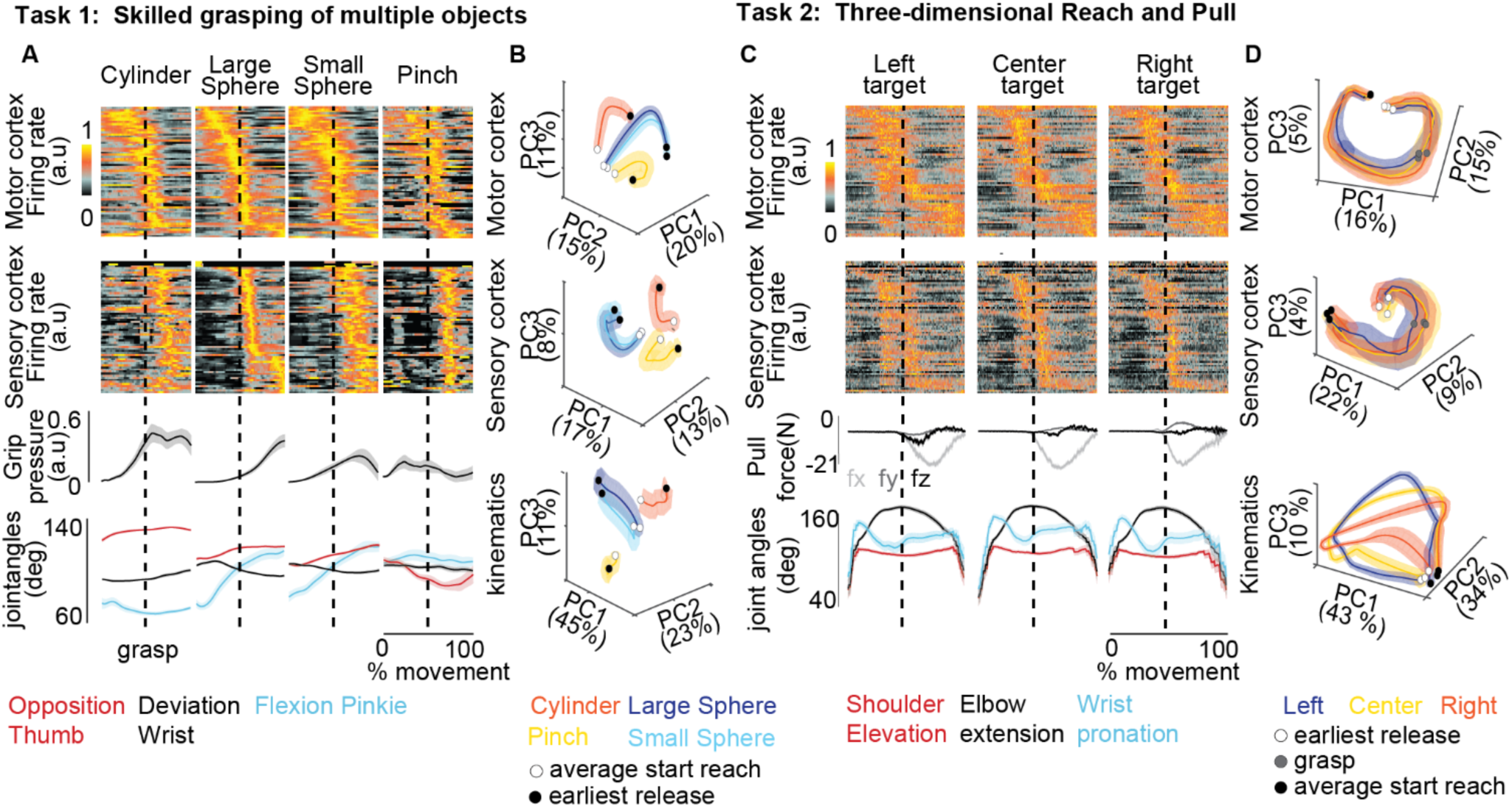
Sensorimotor neural dynamics during unconstrained reaching and pulling. **(A)** Averaged synchronized neural, kinematic and dynamic signals collected during skilled grasping of four different objects (Mk-Jo, n_=_15 trials per object, one recording session). The dashed line identifies the onset of the grasp. Signals were averaged before the grasp over the mean reaching phase duration and after the grasp up to the minimum pulling phase duration. Brain signals were binned over 10 ms time window, normalized for each channel and sorted in ascending order of time of maximal firing rate. The reference condition for aligning neural signals is the large sphere for task 1. **(B)** Principal component analysis of the kinematic, motor and sensory neural components of the movement reveal clustering of the four different types of object geometry along the three first PCs in each the cortical and kinematic PC spaces (Mk-Jo, n_=_15 trials per object, one recording session). **(C)** Averaged synchronized neural, dynamic and kinematic signals during the execution of the task 2 for three different spatial targets. Signals were processed identically to Task 1 (Mk-Cs, n=25 trials per position, one recording session). The reference condition for aligning neural signals is the central target for task 2 **(D)** Principal component analysis of the kinematic, motor and sensory neural components of the movement for the three different object positions (Mk-Cs, n=25 trials per position, one recording session) show overlapping trajectories for the three targets in the neural PC space and dissociated trajectories in the kinematics PC space.

We then identified a neural manifold (Gallego et al., 2017) from the multi-unit activity of each brain area and compared the trajectories to those of the kinematics (Figure 6A and C). In Task 1, both M1 and S1 showed smooth curves that segregate for different object shapes. The trajectories corresponding to similar objects were found to be very close in the PC space for both kinematic and neural features. In Task 2 instead, trajectories in the kinematic manifold showed distinct paths for different positions in space (Figure 6 B, D), while neural manifolds displayed very similar trajectories for different positions suggesting a common neural basis for reaching and pulling in both motor and sensory areas.

#### Single unit encoding of multimodal motor and sensory information

We then exploited the capabilities of our framework to dissect the role of kinematic, kinetic or sensory events on the firing patterns of single units. We manually sorted M1 and S1 recordings to identify well-isolated single units. During the task, firing rates in M1 were typically higher than in S1 (Figure 7A). By design, the pulling force applied by the monkey correlated with the negative (towards the body) hand velocity of the pull (Figure 7B). We first asked whether this interaction between limb kinematics and object dynamics could be observed at the single neuron level. We then plotted the firing patterns of single neurons against force and velocity variables and observed diverse and complex tuning for hand velocity and pulling force in both M1 (Figure 7C) and S1 (Figure 7D). While some cells showed apparent tuning for specific features (Figure 7C top and Figure 7D top likely modulate only with pulling force) others showed a more complex interaction of these features (e.g., Figure 7D, bottom, modulates with both hand velocity and pulling force).

**Figure 7.**
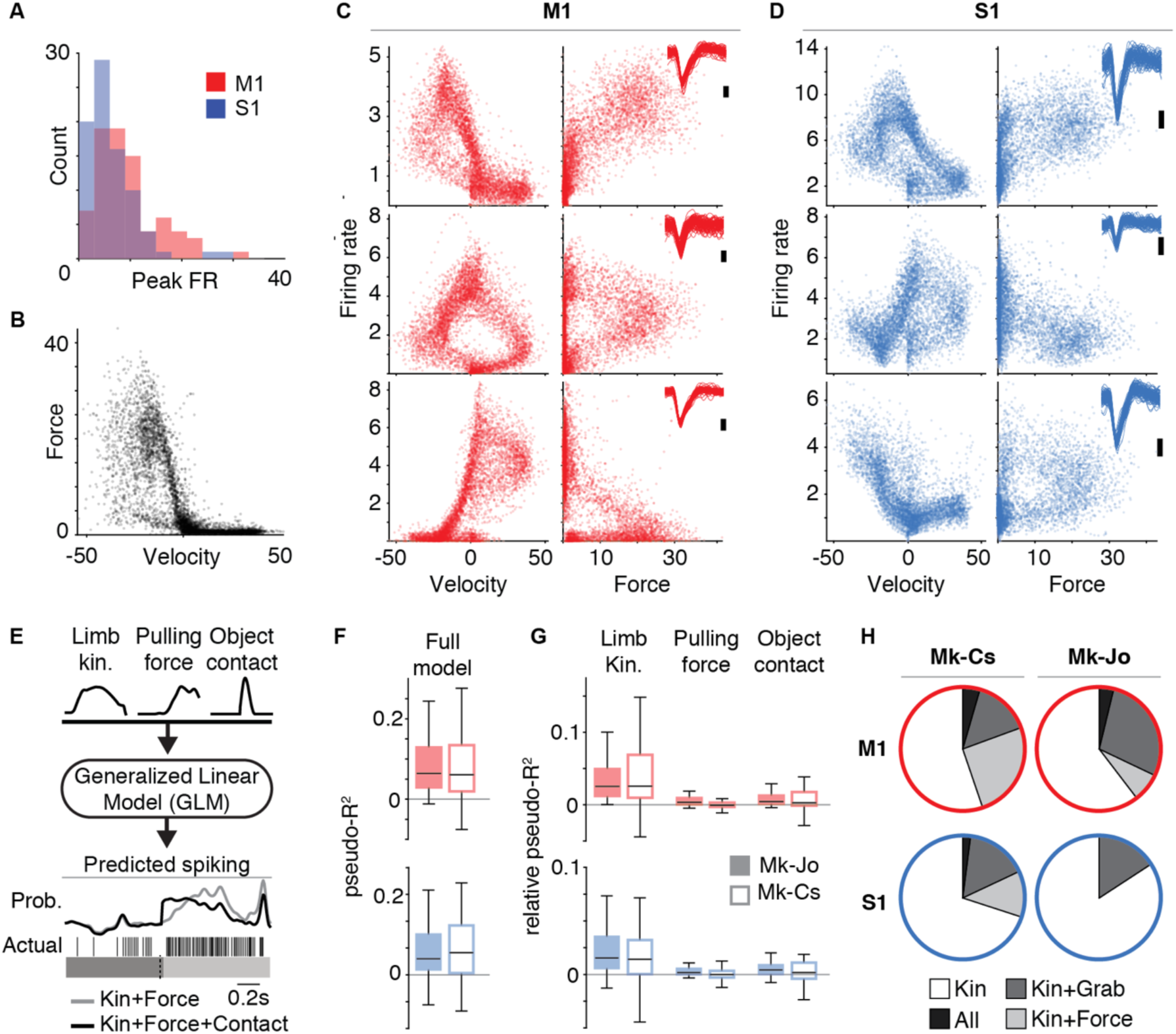
Single unit encoding of multimodal motor and sensory information. **(A)** Each histogram shows the peak firing rate across all trials (one session per animal) for the populations of sorted neurons from M1 (red) and S1 (blue). On average, M1 firing rates were slightly higher than S1 (9.5 Hz compared to 6.4 Hz; p < 0.001, student’s t-test). **(B)** Hand velocity against pulling force for all trials from Mk-Cs. Each dot represents a single time point (sampled at 10ms intervals). Since the pulling phase was loaded, the pulling force strongly correlates with negative hand velocity. **(C)** Example tuning for three M1 cells plotting firing rate against the hand velocity (left column) and pulling for (right column). Insets show 100 randomly drawn spike waveforms (scale bar indicates 100 uV). Due to the interplay between kinematics and force during the object interaction, we observed very complex tuning for the different cells. The top cell, for example, correlates with pulling force, whereas the middle cell appears to correlate with both force and kinematics. **(D)** Three example cells from S1, presented as in Panel C. **(E)** We used GLMs to assess whether neural activity is explained by limb kinematics, pulling force, or object contact events. Example predictions for one cell is shown comparing a model with kinematics and force (gray) against the full model (black). The black model better captures the burst of spikes following object contact. **(F)** The distribution of pseudo-R2 values for the full model across all M1 and S1 cells in Mk-Cs (solid) and Mk-Jo (hollow). **(G)** The relative pseudo-R2 metric captures the contribution of each parameter to the full model fit. Most cells were predominantly explained by limb movements, but there remains a substantial effect of pulling force and object contact. **(H)** The percentage of cells that were significantly described (p < 0.05, bootstrap test; see Methods) by each of the parameters. All cells were significantly predicted by kinematics, though many also were significantly explained by object contact and pulling force.

We next sought to quantify the influence of these behavioural covariates on each spike train using an encoding model of neural activity. We constructed Generalized Linear Models (GLMs) (Perich et al., 2018; Pillow et al., 2008) to predict the spiking activity of the individual neurons based on numerous behavioural and environment signals including limb kinematics, pulling force, and object contact events reflecting the possibility for cutaneous sensory input (Figure 7E) at object contact. The GLMs predicted the probability of observing an individual spike train. We found that the majority of cells in both M1 and S1 could be significantly predicted using this model (Figure 7F), with similar performance for both areas. By computing a relative pseudo-R2 metric, which compares the performance of the full model to a reduced model which omits specific variables (see Methods), we quantified the unique contribution of each specific set of variables on the model performance (Figure 7G). We found limb kinematics to be the dominant explanatory variable, reflecting the gross modulation of neural activity throughout the reach. Yet, a significant portion of neural spiking could also be explained by the three-dimensional pulling force or the time of object contact. For both monkeys, approximately 16% of S1 neurons were significantly explained by the precise object contact independently of other variables (Figure 7H). Across the population of neurons recorded from the M1 arm area Mk-Cs, 24% of neurons were significantly explained by pulling force, yet we found very little force information in the hand-area M1 neurons recorded from Mk-Jo.

#### Discrete and Continuous decoding of movement kinematics

Finally, we sought to demonstrate the relevance of our framework to neural engineering applications. For this, we evaluated the ability to decode discrete and continuous features of natural movement using tools widely employed in studies of Brain Computer Interfaces (Capogrosso et al., 2016; Ethier et al., 2012; Fitzsimmons et al., 2009; Hu et al., 2018; Nishimura et al., 2013; Vargas-Irwin et al., 2010; Velliste et al., 2008).

We developed a linear discriminant analysis (LDA) decoder that calculated the probability of reaching onset and grasping events using cortical signals from primary motor areas. The decoder accurately predicted these events over extended periods of behaviour in both monkeys (Figure 8A), with accuracy of up to 99% for Mk-Jo and 93% for Mk-Cs. The median temporal precision was found to be close to zero (median difference: –11.5 ms for reaching onset and 9.5 ms for grasping in Mk-Jo, and −4ms for both events in Mk-Cs).

**Figure 8.**
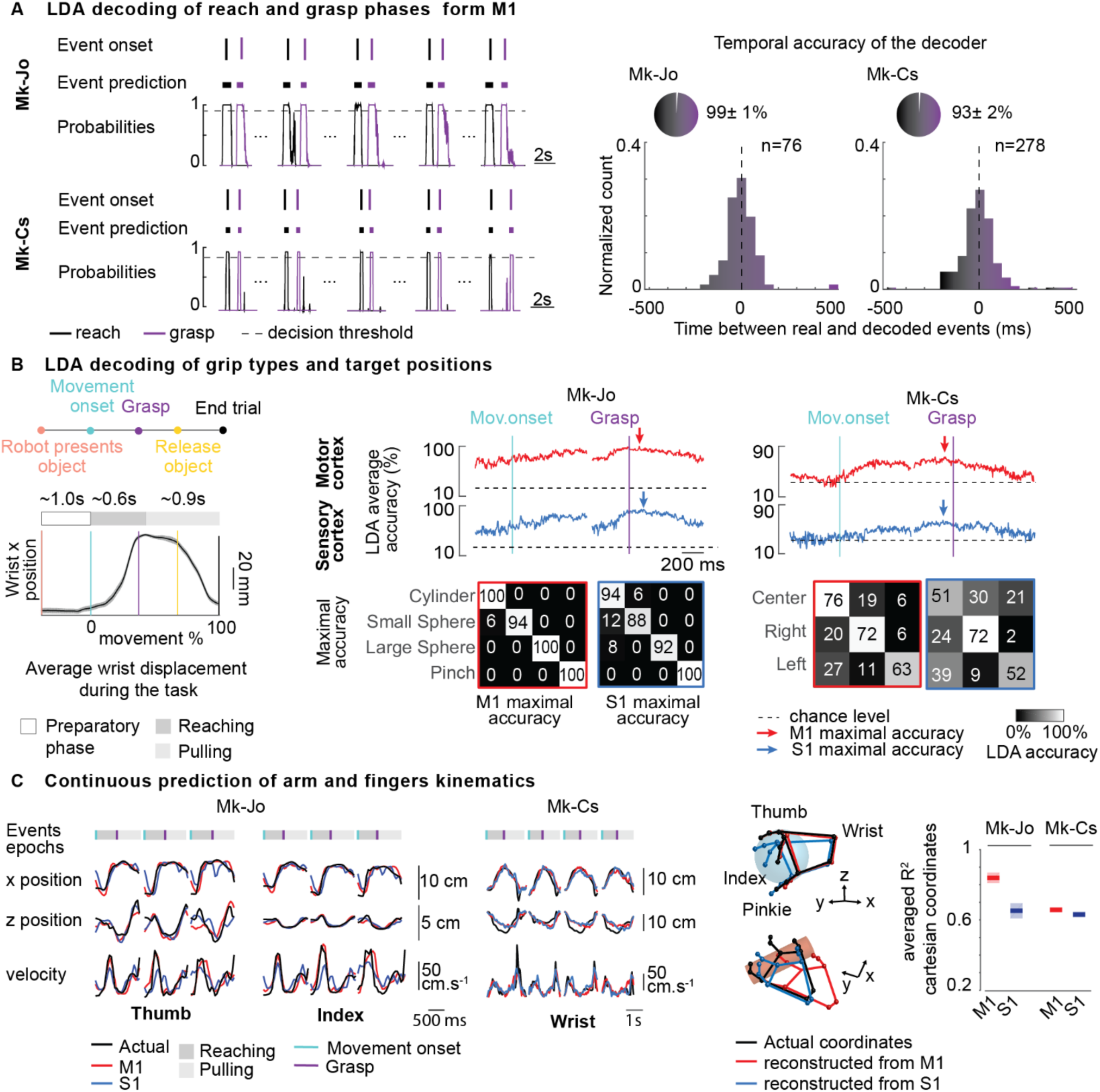
Discrete and Continuous decoding of movement dynamics. **(A)** LDA decoding of reach-onset and grasp-onset events from M1 multiunit activity in both monkeys. On the left, four (Mk-Jo) and five (Mk-Cs) successive reach, grasp and pull trials are presented along with the actual event onset identified using video recordings, the detected motor states and the probability of reach-onset and grasp-onset motor states (in black and purple). On the right, histograms show the distribution of the temporal differences between the actual occurrence of reach-onset and grasp-onset events and the decoded occurrence of these motor states for all the recording sessions in the two monkeys (n=76 reach and grasp events for M-Jo, one recording session, and n=278 reach and grasp events for Mk-Cs, one recording session). **(B)** LDA decoding of grip types and target positions. On the left, we represented the trial structure as well as the average wrist displacement during the task (n=48 trials for Mk-Jo, one recording session). The wrist position does not vary before the movement onset event identified via video recordings. The average reaching phase duration was 650 ms, while the average pulling phase lasted for about 900 ms On the right, we show the average performance of the offline LDA decoder for each object shapes and each target position in space using multiunit activity recorded from M1 and S1 areas (n=15 trials per objects for Mk-Jo, one recording session and n=25 trials per position for Mk-Cs, one recording session). Confusion matrices report accuracies obtained for the best time window for M1 and S1 signals. **(C)** Continuous prediction of arm and fingers kinematics using Kalman filtering. Multiunit activity from M1 and S1 areas were used to predict fingers and arm 3D coordinates during the whole reach-and-pull movement. On the left, time traces show prediction of the thumb, index and wrist x, y positions and velocity using neural population activity from M1 and S1 arrays in Mk-Jo and Mk-Cs. Three-dimensional static hand postures were reconstructed by predicting x, y and z coordinates of each fingers and wrist joints in Mk-Jo for the small spherical grip and the cylindrical grip (n=1 trial per object). On the right, performances of the continuous decoders are computed separately for M1 and S1 in both monkeys (cross validation k-folds with k=48 for Mk-Jo, one recording session and k=134 for Mk-Cs, one recording session).

We then investigated the LDA-based decoding power of object type and direction of reaching in both motor and sensory areas. We repeated the LDA analysis at sliding time windows spanning the entire reach-and-grasp movement. Figure 8B shows average confusion matrices obtained for the best time windows. Both M1 and S1 information could be used to discriminate between different grasps during the grasping phase in Mk-Jo (average maximal performance was 99% for M1 and 94% for S1). Interestingly, multiunit activity from M1 and S1 could be used to differentiate between objects more than 300 ms before movement onset. Yet, the exact movement direction in space could not be reliably predicted before the movement onset in Mk-Cs. Figure 8B shows that the maximal decoding accuracy was reached around 100 ms before the grasp (average maximal performance was 70% for M1 and 58% for S1).

We finally investigated the possibility to decode continuous arm and fingers kinematics. By fitting a standard Kalman filter (Wu et al., 2006) to each 3D coordinates of the arm and hand joints, we managed to reconstruct time-varying joint coordinates for most of the finger joints, obtaining a median performance of 0.83 and 0.65 for M1 and S1 areas respectively in Mk-Jo (Figure 8C). Three-dimensional reconstruction of the wrist and fingers position based on decoded kinematic features is shown in Figure 8C for both spherical and cylindrical grips. Hand and arm joint kinematics could also be decoded from Mk-Cs with a median performance of 0.68 and 0.67 for M1 and S1 area respectively. For both monkeys, M1 decoder outperformed S1 decoder.

## DISCUSSION

Here we presented the implementation of a robotic platform for the study of sensorimotor processes underlying upper limb movements in monkeys. We validated this platform by implementing behavioural tasks in monkeys involving natural, three-dimensional reaching, grasping and object manipulation. We discuss our developments and the functionality of our robotic platform in light of three applications: 1) the study of arm/hand kinematics, 2) neural correlates of natural upper limb movements 3) application in Brain Computer Interfaces.

### A versatile framework for multimodal characterization of three-dimensional arm movements

In typical neurophysiological experiments with monkeys, experimenters constrain the behaviour of monkeys to separate kinematic analysis from the study of dynamic force interactions with the world. Reach to grasp is often studied using static objects (Brochier et al., 2004; Ethier et al., 2012; Okorokova et al., 2019; Santello et al., 1998; Schaffelhofer and Scherberger, 2016; Vargas-Irwin et al., 2010) or constrained manipulandum (de Haan et al., 2018; London and Miller, 2013; Omrani et al., 2016), while the production of forces is typically investigated under isometric/unidimensional conditions (Ethier et al., 2012; Moritz et al., 2007; Nishimura et al., 2013) or in planar tasks (de Haan et al., 2018; London and Miller, 2013; Omrani et al., 2016). While these constraints are effective to dissect the role of specific neuronal-subpopulation in the control of dynamic movement variables, even simple natural movements present an extraordinary complex patchwork of unconstrained, precision and force-controlled movement actions. Our robotic framework allows the characterization of these natural behaviours while retaining the ability to integrate multimodal recordings of kinetic and kinematic signals.

We and devised a platform in which subjects can interact freely with a robotic arm in a large three-dimensional workspace (Figure S1 C). We showed that interaction with the robot during upper-limb movements was safe and robust and that it did not impose any direct physical constraint on the arm, therefore promoting natural and smooth trajectories (Figure 3F). In addition, we instrumented our platform to provide simultaneous recordings of pulling and contact forces. Our robot control strategy can accommodate for simple and straightforward definition of motor tasks and it allows the user to easily adjust robot behaviour. The control software package and libraries are made freely accessible and open source on Zenodo (<u>10.5281/zenodo.3234138)</u>. We finally synchronized kinematics and cortical recordings to both robot behaviour and interaction forces measurements to obtain a rich multimodal panel of sensory and motor features characterizing arm and hand movements.

This framework provides with unprecedented means to address scientific questions that require the study of unconstrained and natural movements in monkeys such as the characterization of motor recovery in pre-clinical studies. It can potentially be used to design complex tasks such as precise control of grip pressure (Ethier et al., 2012), maneuvering of 3D manipulandum, or motion under force-field perturbation. In addition, our platform could serve rehabilitation purposes (Spalletti et al., 2017; Spalletti et al., 2014). For example, deficits and recovery in kinematics or force outputs could be characterized in animal models of disease.

### Applications in the study of three-dimensional movement dynamics

Characterization of arm and hand movement dynamics requires the integration of pulling force components, contact forces and kinematics of the joints. In our example, we utilized these features to define a “dynamic” manifold showing the evolution of trajectories in a space that mixes kinematic and forces. We used this tool to study kinematic adaptation to resistance levels and investigate the relationship between kinematics and pulling forces in grasping (Santello et al., 1998). Surprisingly, our analysis revealed that kinematic strategies adapted to resistive loads. This suggests that kinematic synergies may vary when stronger forces are perceived. Our results indicate that during functional movements, upper-limb postures, interaction forces, and joint trajectories are continuously adapted together to reach the desired motor output, and perhaps regulated by both central (Santello et al., 1998) or even simple reflex mechanisms (Weiler et al., 2019).

### Applications in the study of neural sensorimotor processes

We then sought to demonstrate the efficacy and potential of our framework for applications in the study of neural dynamics during natural movements. We reported that neural activity from both motor and somatosensory cortex show clear modulation with kinematic and dynamic movement variables. Both M1 and S1 activity showed complex modulation throughout the whole movement, with clear and smooth trajectories in the neural manifold (Figure 6). This corroborates that somatosensory single cells encode multiple movement components and not only, or mostly, touch-related information (London and Miller, 2013), (Prud’Homme and Kalaska, 1994). The firing patterns of single units in both M1 and S1 showed strong and complex modulation with respect to multiple movement parameters (Figure 7). Our unique set up allowed us to investigate how kinematic, forces and contact events could, in combination or independently, explain neuron firing patterns. The majority of neurons in the somatosensory cortex encoded kinematics variables only. The remaining portion of cells encoded either contact events or force in conjunction with kinematics suggesting that important components of movement dynamics are encoded in somatosensory area 2. Availability of such a complex dataset and integrated kinematic signals opens intriguing possibilities for the study of population dynamics during natural behaviour.

### Applications in neuroengineering and brain computer interfaces

Brain Computer Interfaces have been applied to animal models and humans to control robots (Collinger et al., 2013; Hochberg et al., 2012; Hochberg et al., 2006; Wodlinger et al., 2015) support communication (Gilja et al., 2015; Milekovic et al., 2013b; Vansteensel et al., 2016; Wolpaw et al., 2002) and restore motor control (Ajiboye et al., 2017; Bouton et al., 2016; Ethier et al., 2012; Moritz et al., 2007). However clinical applicability is limited by the fact that these devices are tested in restricted laboratory environments and constrained tasks. Instead, the development of future solutions aiming at the recovery of functional movements requires set-ups that replicate and quantify performances of natural motor tasks. Our framework provides with such an opportunity allowing the execution and quantification of three-dimensional arm and hand movements in a relevant animal model.

For example, we showed that it was possible to decode both reach and grasp events as well as as target shape or position in space during a natural tasks that did not constrain the execution time or the movement trajectory. This type of information could be combined to build simple though robust decoders driving pre-programmed patterns of stimulation of the nervous system in pre-clinical and clinical settings (Capogrosso et al., 2016; Wagner et al., 2018).

Interestingly, even in a task that did not constrain the duration of the movement preparation or execution phase, we were able to predict grasp types from both motor and sensory activity several hundreds of milliseconds before movement onset. We found this result in accordance with several theories around the dynamical interaction and integration of sensory-motor processed during movement, and may even include components such as “efference copy” (Grüsser, 1994).

Finally, we showed that we were able to predict individual fingers kinematics from multiunit activity recorded in motor and somatosensory cortex during whole three-dimensional reaching and grasping movements (Figure 8C). These results confirm the findings of Vaskov and colleagues (Vaskov et al., 2018) on M1-based finger motion decoding and extend the findings of Okorokova and colleagues (Okorokova et al., 2019) to multiunit activity and whole arm and hand movements.

### Conclusion

In summary we have reported and described a platform that can be adapted to study a large variety of tasks that are useful to investigations in motor control and perhaps other related applications. We provided three examples showing the validity of our approach for both basic and applied investigations thus paving the way to more detailed studies investigating sensorimotor processes in both monkeys and humans.

## METHODS

### Animals involved in this study

Two adult female *Macaca fascicularis* monkeys were trained in this study (Mk-Jo 10 years old, 3.6 kg and Mk-Cs 9 years old 4.0 kg). All procedures were carried out in accordance to the Guide for Care and Use of Laboratory Animals (ISBN 0-309-05377-3; 1996) and the principle of the 3Rs. Protocols were approved by local veterinary authorities (authorizations No 2017_03_FR and 2017_04E_FR) including the ethical assessment by the local (cantonal) Survey Committee on Animal Experimentation and acceptance by the Federal Veterinary Office (BVET, Bern, Switzerland).

### Surgical procedures

All the surgical procedures were performed under full anaesthesia induced with midazolam (0.1 mg/kg) and ketamine (10 mg/kg, intramuscular injection) and maintained under continuous intravenous infusion of propofol (5 ml/kg/h) and fentanyl (0.2-1.7 ml/kg/h) using standard aseptic techniques. A certified neurosurgeon (Dr. Jocelyne Bloch, CHUV, Lausanne, Switzerland) supervised all the surgical procedures. We implanted two 64-channel microelectrode arrays (Blackrock Microsystems, 400 µm pitch and electrodes tip lengths 1.5 mm and 1mm for M1 and S1 respectively) Mk-Cs was implanted in M1 and S1 area of the arm, and Mk-Jo was implanted in M1 and S1 area of the hand. Functional areas were identified with electrical stimulation delivered as byphasic pulses on the cortex surface at 3 mA and 300Hz. A 20 mm diameter craniotomy was performed in order to span the brain areas of interest and the dura was incised. Implantation of the arrays was achieved using a pneumatic compressor system (Impactor System, Blackrock Microsystems). The pedestal was fixated to a compliant titanium mesh modelled to fit the skull shape. Surgical and post-operative care procedure are developed in details in (Capogrosso et al., 2018). Data presented in this paper were collected 3 weeks and 9 weeks post-implantation for Mk-Jo and Mk-Cs respectively.

### Behavioural tasks

We built a custom primate chair (Supplementary Figure S1) in which only the neck of the animal was fixed with a metallic collar to permit a wide range of voluntary arm and hand movements in the three-dimensional space. Two Plexiglas plates were used to restrain the access to the head and a third plate placed around the womb of the animal served for placing a resting bar and elastic bands to immobilize the right hand. Monkeys were trained to maintain a resting position between trials and place the left hand on a metallic bar a few centimetres in the front, at chest level. A trial started when the robot presented a graspable target in front of the animal, at a distance of approximatively 20 cm. As soon as the “go” cue was played (1 sec duration sound), monkeys reached for the target, grasped it, and pulled it towards themselves. Once the robot end effector crossed a pre-set virtual spatial threshold (8 cm), a clicker sound played by the experimenter indicated the success of the trial encouraging the animal to release the grip, return to the resting bar and get a food reward. The robot returned to its vertical home position at the end of each trial. Cueing signals during the task were implemented as follow: a go cue was implemented as a high pitch sound coupled to a green light that was played when the robot was in position and reached the impedance control mode; a success cue consisted in a low pitch sound when the monkey crossed the spatial threshold by pulling on the robot. Sounds were played from the application controlling the robot while the light cue was triggered through our synchronisation module (Arduino Due, Arduino, Italy).

In Task 1, Mk-Jo performed a similar task in which objects of different size and shape were presented. A cylinder (length 8 cm; diameter 1.5 cm), a small sphere (diameter 1.5 cm) and a large sphere (diameter 3 cm) were used to induce cylindrical, small and wide spherical grips respectively. CAD design of these objects are available at (<u>10.5281/zenodo.3234138)</u>.

. Each of these targets were presented with various levels of resistance applied by the robot impedance controller (joint stiffness 200, 400 and 600 N/m). In addition, a small triangular pinch-like object (base 2 cm; height 1.5 cm) was used to prompt a lateral precision grip and was presented with the lowest level of resistance. Each session consisted in 12 to 20 trials per object per level of resistance. Two to four objects were presented during a single session.

In Task 2, Mk-Cs was trained to perform 3-dimensional centered-out task where the small spherical object was alternatively and randomly presented in three horizontal positions in the sagittal plane, center, left and right. The object was placed at −40, 0 and 60 mm along the y-axis for the left, center and right position respectively. The z distance was fixed for all conditions at approximately 180 mm above the animal seating height. Each session consisted in approximately 25 trials per position.

### Robotic arm control

The control architecture of the robotic framework consists in a finite state machine including five states. At the beginning of a session the robot lies in *home* position (Figure 2, FS1), in which all joint coordinates equal to 0°, resulting in a straight and vertical robotic arm configuration (supplementary Figure S1C). The robotic arm maintains its position until the user induces the start of a new trial by pressing a remote button. The pressure on the button brings the robot in a position control phase (Figure 2, FS2), during which it reaches for the target position. Once the sum of the errors in the positioning of all the joints decreases to a value of 0.01 radians, the robot is considered in target position and the control mode switched to an impedance control mode (Figure2, FS3). During this finite state, the robot behaves as a mass-spring-damper system, trying to keep the target position while opposing a certain degree of resistance to any applied force. In our particular impedance control strategy, we facilitated a smooth motion of the end effector along the x-axis, by imposing the higher stiffness and damping parameters along the y and z direction. The monkey was trained to grasp a flexible and sensorized object anchored to the robot end effector, and pull it until crosses a user defined position along the x-axis. Upon success, the robotic arm quickly switches to FS4 and FS5 during which the robot automatically updates the next desired target position and regains the home position. Upon failure, target position is not updated and the robot directly reverts to the home position. After three failed trials the robot switches to another target position regardless of the behavioural outcome.

### Multimodal recordings

#### Kinematic recordings

Three-dimensional spatial coordinates of arm and hand joints during upper limb movements were acquired using a 14-camera motion tracking system (Vicon Motion Systems, Oxford, UK) at a 100 Hz-frame rate. The video system tracked the Cartesian position of up to 15 infrared reflective markers (6 to 9 mm in diameter each, Vicon Motion Systems, Oxford, UK). For each monkey, one marker was placed directly below the shoulder, three on the elbow (proximal, medial and distal) and two were placed on the wrist (lateral and medial) using elastic bands. For Mk-Jo, nine additional markers were positioned on the back of a customized viscose glove, on the metacarpal (MCP), proximal (PIP) and distal phalanxes (DIP) joints of the thumb, index and little finger (Supplementary Figure S2A). A model of each subject’s marker placement was calibrated in Vicon’s Nexus software.

#### Pulling force recordings

The interaction force, measured as the force applied at the robotic joints, was sampled at 500 Hz and stored in a text file along with triggers marking the beginning (go cue) and the end (spatial threshold crossing) of each trial and the spatial position of the target object. Values were also streamed as digital output to the BlackRock recording system using the single-board microcontroller. The pressure sensor was fixed to the robot end effector and analog inputs and outputs were passed through a custom-made cable plugged at the base of the robot. The analog pressure signal was sampled at 1000 Hz in the VICON system and synchronized with kinematics triggers marking the start (go cue) and the end (threshold crossing) of each trial.

#### Electrophysiology recordings

Neural signals were acquired in the BlackRock system using the Cereplex-E headstage with a sampling frequency of 30 kHz. Multiunit activity was extracted offline for each of the 128 channels through filtering (250Hz-5kHz) and spiking events were extracted through threshold crossing (Capogrosso et al., 2016). Specifically, a spiking event was defined on each channel (2×64 in total) if the signal exceeded 3.0–3.5 times its root-mean-square value calculated over a period of 5 s. Artefacts removal was achieved by eliminating all the spikes occurring within a time window of 0.5 ms after a spike event in at least 30 channels. Firing rate was computed for each channel as the number of spikes detected over a sliding window of 100 ms.

### System performances characterization

We evaluated safety and usability of the robotic framework by performing an experiment on a naive human subject (see section Results). We continuously recorded the end effector three-dimensional position, together with the three-dimensional pulling force. We computed both a *precision error,* a *drop error* (Figure 3A). The precision error was computed as the distance between the targeted end effector position and the reached endpoint position, therefore illustrating the reproducibility of the robotic arm across trials. The drop error was computed as the three-dimensional drift of the end effector from the target position during the first 500 ms after placement, illustrating the stability of the robotic arm in holding the object in the target position. Subsequently, we verified the compliance of the robot movement upon interaction with a monkey previously trained to perform Task 2. We recorded three-dimensional coordinates of the joints and then computed a *trajectory smoothness index* (Hans-Leo Teulings et al., 1997)computed as:

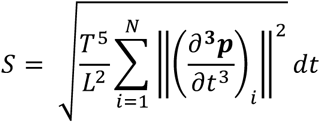

where ***p*** is the end effector position, N is the number of samples considered for the measure, *t* is the time, T and L represents the total duration of the trajectory in seconds an the length of the trajectory in m respectively. We computed a separated smoothness index over the reach and the pull.

All the grip pressure sensor-coupled objects were characterized using a computer-controlled compression system (Zwick/Roell 1KN D0728165) to derive the internal pressure of each object as a function of the applied force in Netwon. Calibration curves were acquired by applying dynamically load ranging from 0 to 25 N for the large sphere, 0 to 20 N for the cylinder and small sphere and 0 to 10 N for the pinch. Fitting curves were computed post-hoc using standard Matlab (The Mathworks fitting tools and were used to correct for the grip pressure prior to data analysis.

### Pressure sensor implementation

The pressure sensor circuit converted air pressure into a voltage measurement using strain-gages (1620 measurement SPECIALTIES^TM^) directly connected to the front-end amplifier (FEA - LMP90100) of a full-Wheatstone bridge circuit. The footprint of the electronic circuit was maximally reduced in order to integrate it to the robot (diameter 3.2 cm). The embedded circuit converted the strain-gages (1620 measurement SPECIALTIES^TM^) signal into an analog output signal bounded between 0 and 4.1V. The electronic assembly comprised a microcontroller (pic24FV08KM101) for the control of the AFE and the digital-to-analog converter (DAC8551) via a serial peripheral interface (SPI) protocol. A front-end amplifier performed the 16 bits digital-to-analog conversion, the amplification and the noise reduction of the signal. A digital to analog converted (DAC) converted the processed signal. A 5V regulator (EG113NA-5) adjusted the input power supply from 6V to 9V and a second regulator (LM4140ACM-4.1) outputted a precise reference voltage of 4.1V used for signal conversions. Three LEDs indicated the state of the system and three input switches were used to impose gains ranging from 1 to 64. The in-circuit serial programming (ISCP) connector served to re-program the microcontroller. The system was also equipped from an ON/OFF button, a reset button and a power supply connector.

The sensor and the electronic circuit were placed in an inox case and coupling of the instrumented objects to the sensor was achieved with a custom-made excavated screw, to allow rapid and easy adjustment of the object. The assembly was mechanically and electrically coupled to the flange of the robotic arm.

All the sensitive elements were characterized using a computer-controlled compression system (Zwick/Roell 1KN D0728165) to derive the internal pressure of each object as a function of the applied force.

### Analysis of arm and hand dynamics

Post processing of motion capture data was performed to ensure that all joint markers were labelled correctly. We converted the three-dimensional marker position data to rotational DoFs. For Mk-Jo, we computed 12 joint angles (see Figure S2): index finger PIP flexion/extension, pinkie finger PIP flexion/extension, thumb finger PIP flexion/extension, index finger MCP flexion/extension, pinkie finger MCP flexion/extension, thumb finger MCP flexion/extension, index finger abduction/adduction, pinkie finger abduction/adduction, thumb finger abduction/adduction, thumb opposition, wrist flexion, wrist ulnar deviation. For Mk-Cs, we computed 3 joint angles: shoulder adduction, elbow flexion/extension angle and wrist pronation/supination. Interaction force signals were synchronized post-hoc to kinematics signals based on triggers marking the start (go cue) and end (threshold crossing) of each trial. For further analysis of arm and hand dynamics during natural reaching and grasping, signals were averaged over each condition. Kinematics, force and grip pressure signals were low-pass filtered at 10 Hz.

Principal Component Analysis (PCA) was performed on joint angle kinematics and kinetic variables to identify which dynamic features accounted for most of the variance in the data. We reconstructed the Y x N_df_ matrix where N_df_ is the number of joint angles and *Y* = [*Y*_1_, *Y*_2_,…, *Y_N_*] is the concatenated vector of all time points *t* = 1,…, *N*. Average kinematic, force and pressure variables were normalized and mean-centered before singular values decomposition. The dynamic data projected along the first three eigenvectors in the PC space were averaged over each condition and smoothed using a moving average filter before plotting.

### Analysis of kinematic synergies

We used PCA (as described in (Santello et al., 1998) to identify kinematic synergies as orthogonal axes of maximal correlated variance in the 3D joint coordinates of the hand. Briefly, the x, y and z joints Cartesian coordinates were first normalized and mean-centered. The principal components (PCs) were then computed from the eigenvalues and eigenvectors of the matrix of the covariance coefficients between each of the joint coordinates waveforms. As each of the 13 markers was represented by three coordinates the PCA was computed over 39 waveforms. The first PC accounted for at least 70% of the variance. We represented the hand posture corresponding to the first synergy by plotting the 39 coefficients corresponding to the first PC in 3D (Mason et al., 2001) and overlaid a sketch of a monkey hand for the sake of representation. The positions of the ring finger and middle finger were not measured in our experiment but were represented in in a realistic inferred position with respect to the thumb index and pinkie to help visualize the hand posture in supplementary Figure 2D.

### Multiunit activity analysis during skilled grasping and 3D reaching

In Figure 6, 2D activation plots displayed the average firing rate computed over 100 ms window. The firing rates were normalized for each channel and sorted so that channels reaching their peak (maximal) activity first would be displayed on top. For Mk-Jo, channels were aligned according to their peak activation during large spherical grasp while for Mk-Cs, channels were aligned according to their peak activation during reaching towards the central target.

PCA analysis was conducted on neural firing rate and kinematic data ranging from movement onset to the end of the pulling phase. In Mk-Jo, movement onset happened in average 660ms before the grasp and the pulling phase lasted at least 350 ms. In Mk-Cs, movement onset happened in average 910 ms before the grasp and the pulling phase lasted at least 740 ms. Events marking movement onset, grasp onset and release of the object were identified using video recordings. Neural features for M1 and S1 consisted in the firing rates computed over all the 64 channels of each array. Kinematic features corresponded to the same kinematic features described in Figure 5C and F for Mk-Jo and Mk-Cs respectively. PCs were computed using at least 15 trials per object and 25 trials per position.

### Neural encoding analysis

We manually spike-sorted the recordings from each electrode implanted in M1 and S1. We computed the average firing rate across the entire reach for each neuron to compare the two brain areas (Figure 7a). Additionally, we inspected the tuning for each parameter by plotting the firing rate in each bin against the hand velocity and the magnitude of pulling force recorded at each time (Figure 7b).

We then constructed encoding models to predict the spiking of each neuron using Generalized Linear Models (GLMs), adapting an analysis previously described by (Lawlor et al., 2018; Perich et al., 2018). First, we counted the number of spikes in non-overlapping 10ms bins. In brief, GLMs generalize the idea of multilinear regression for non-Gaussian statistical distributions using a nonlinear link function. The neurons were assumed to have Poisson statistics, thus we used an exponential link function (Lawlor et al., 2018). As inputs to the GLMs we provided: 1) full-limb kinematics, including the three-dimensional velocities and accelerations of the shoulder, elbow, and wrist; 2) three-dimensional pulling force applied to the robot by the monkey; 3) the time of the grab event, to capture either motor commands related to hand shaping or sensory feedback from the object contact. For the third input, the grab event was convolved with three raised cosine basis functions (Pillow et al., 2008) spaced evenly up to 300 ms in the past or future for M1 and S1, respectively.

We quantified the performance of the GLMs using a pseudo-R^2^ metric, which generalizes the notion of variance explained for the Poisson statistics of the model (Lawlor et al., 2018; Perich et al., 2018). This metric compares the log-likelihood of the tested model fit against a simpler model.

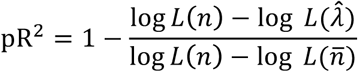

Typically, the simpler model 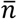 is a mean-fit to the data. However, this formulation also allows us to test the relative contributions of different parameters. We compared the full model with all three types of inputs described above to a reduced model 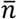 where just one of the inputs is omitted. This *relative pseudo-R^2^*metric provides insights into how well specific parameters, such as object contact, helps to explain neural activity. We assessed significance for all of these using a Monte Carlo simulation resampling across the available trials. A model fit or parameter was assumed to be significant if the 95% confidence intervals on the parameter fit were greater than zero.

### Detection of movement onset and object grasp from the sensorimotor neural activity

We implemented an approach based on a multiclass regularized linear discriminant analysis algorithm (mrLDA) to detect moments of movement onset and object grasp from the continuous neural recordings from either the motor or somatosensory cortices (Capogrosso et al., 2016; Milekovic et al., 2013a; Milekovic et al., 2018). We synchronized the multiunit spike activity with the movement onset and object grasp events identified using video recordings. The performance of decoders was evaluated using five-fold cross-validation(Hastie et al., 2009) - we divided the dataset in five parts of similar lengths, used four of these parts as the training set for decoder calibration and remaining part as the testing set to evaluate the decoder performance. This calibration involves a procedure to select parameters of the mrLDA algorithm. Specifically, movement onset and object grasp events, *mo* and *og*, were used to derive the respective classes of neural features, *C_mo_*and *C_og_*:

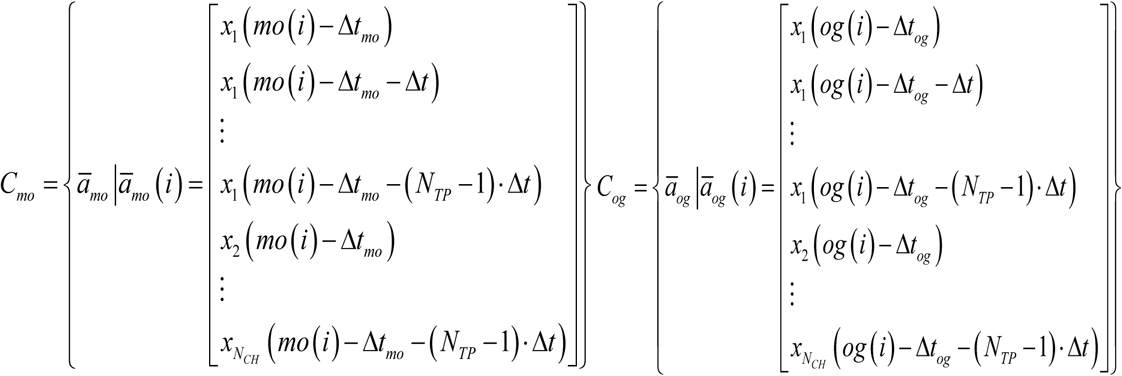

where 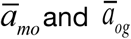 are feature vector members of classes *C_mo_* and *C_og_*, respectively; *N_TP_*is the number of multiunit spike rate measurements taken from the same neural channel; *Δt* is the temporal difference between the two consecutive spike rate measurements taken from the same neural channel; *N_CH_*=64 is the number of neural channels; and *Δt_mo_* and *Δt_og_* are the temporal offsets for each type of event. *N_TP_, N_TP_ ·Δt* (history used to sample neural features)*, Δt_mo_* and *Δt_og_* were used as decoder parameters - their values were selected from a following list of values: *N_TP_*: 3 and 5; *N_TP_ ·Δt*: 0.3s and 0.5s; *Δt_mo_* and *Δt_og_*: any value from - 200ms to 200ms with 0.5ms steps. We formed another class *C_OTHER_*representing states other than *mo* and *og* events:

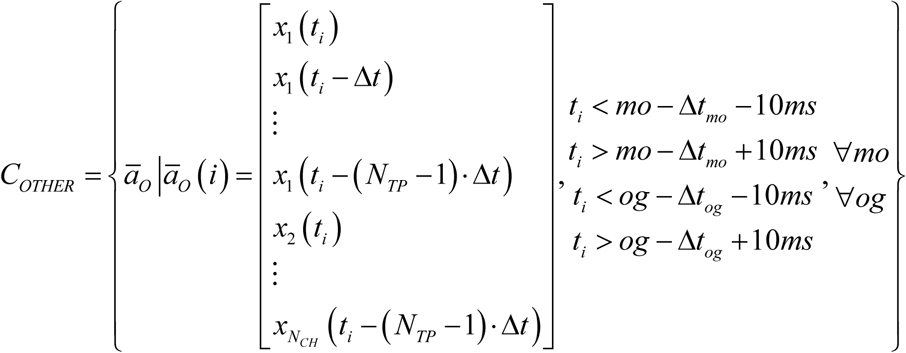

where *t_i_* are all times at least 10ms away from all *mo* and *og* events. We then calibrated a set of mrLDA decoding models using *C_mo_*, *C_og_* and *C_OTHER_* and all possible combinations of parameter values(Capogrosso et al., 2016; Milekovic et al., 2013a; Milekovic et al., 2018). We used the mrLDA regularization parameter as an additional parameter with values of 0, 0.001, 0.1, 0.3, 0.5, 0.7, 0.9 and 0.99. The performance of each of the models was validated using four-fold crossvalidation on the training dataset as follows. A model was calibrated on three quarters of the training dataset and tested on the remaining part of the training dataset. For each time point of this remaining part, the mrLDA model calculated the probability of observing neural activity belonging to *C_mo_* and *C_og_*classes. When one of these two probabilities crossed a threshold of 0.9, the decoder detected movement onset or object grasp, *mo_DET_* or *og_DET_*, respectively. To reproduce the sparsity of these events, we ignored the probability values of the detected event for 1s after the detection. We pooled the time series of actual and detected events across all four folds and calculated normalized mutual information using a tolerance window of 200ms (Capogrosso et al., 2016; Milekovic et al., 2013a; Milekovic et al., 2018). We then calibrated another decoding model on the complete testing part of the dataset using the parameter values that resulted with the maximum normalized mutual information. This decoder was then used on the testing part of the dataset to detect the *mo_DET_* or *og_DET_* events. We again pooled actual and detected events across all five testing folds and measured the decoding performance using temporal accuracy – the ratio of actual events that have one and only one event of the same type within the 200ms neighbourhood. We estimated the standard error of the temporal accuracy using bootstrapping with 10 000 resamples(Moore et al., 2009).

### Time-resolved classification of grasp types and trajectories

We classified grasp types (Mk-Jo) and reach target locations (Mk-Cs) using mrLDA from the neural recordings from either the motor or somatosensory cortex (Milekovic et al., 2015). We synchronized the multiunit spike activity with the movement onset and object grasp events and generated trials – data periods lasting from 1s before to 1s after each event. We measured the classification performance using decoding accuracy evaluated using leave-one-out validation (Hastie et al., 2009) – in each validation fold, one trial is selected as a test dataset and all other trials are used as a training dataset to calibrate a mrLDA model. This calibration involves a procedure to select the regularization parameter of the mrLDA model from the values of 0, 0.001, 0.1, 0.3, 0.5, 0.7, 0.9 and 0.99. The training dataset is used to derive the classes of neural features, *C_1_(e,t)*, …, *C_N_(e,t)* (N=4 for four objects in Mk-Jo, and N=3 object positions for Mk-Cs):

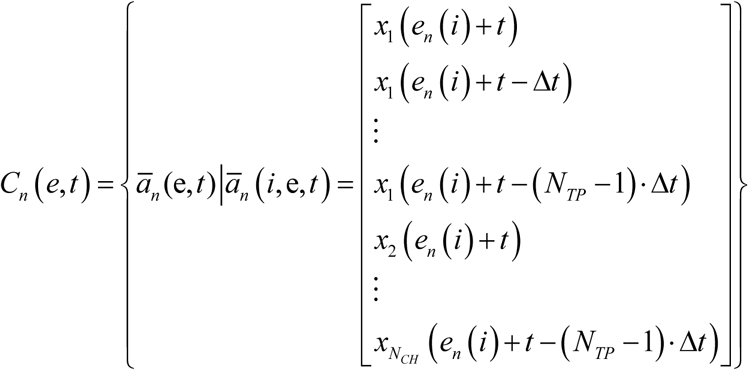

where *e* are movement onset or object grasp events; *t* is time latency (values from −1s to 1s in steps of 5ms) with respect to these events; 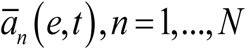 are feature vector members of *C_1_(e,t)* to *C_N_(e,t)*, respectively; *N_TP_*=3 is the number of multiunit spike rate measurements taken from the same neural channel; *Δt*=150ms is the temporal difference between the two consecutive spike rate measurements taken from the same neural channel; *N_CH_*=64 is the number of neural channels; and *Δt_mo_* and *Δt_og_*are the temporal offsets for each type of event. For each time latency *t* and event type *e*, we calibrated a set of mrLDA classifiers using *C_1_(e,t)*, …, *C_N_(e,t)* classes and all possible combinations of the regularization parameter. The performance of each of the models was evaluated using leave-one-out validation on the training dataset. We then calibrated a mrLDA classifier on the complete testing dataset using the regularization coefficient that resulted with the maximum decoding accuracy. We pooled the classification outcomes across all folds and measured the decoding accuracy. This procedure resulted in a confusion matrix and decoding accuracy value for every value of the latency *t* around both movement onset and object grasp events. We estimated the standard error of the decoding accuracy using bootstrapping with 10 000 resamples (Moore et al., 2009).

### Kalman filter decoding

We assessed continuous predictions of limb kinematics using a Kalman Filter (Wu et al., 2006). The Kalman Filter provides a probabilistic framework to predict the state of the limb during the reach and grasp task based on the neural recordings. The output of the filter was the state of the limb. For Mk-Cs, this comprised the position and speed of the elbow and wrist. For Mk-Jo, it comprised the position and speed of the thumb, index and pinkie finger joints (distal phalanx, PIP joint and MCP joint), as well as of the wrist, elbow and shoulder joints. We computed the limb kinematics and the instantaneous firing rate of each neuron at 50ms intervals. We used the multiunit firing rate obtained from thresholding on M1 and S1 arrays as inputs to the decoder. The neural signals were shifted relative to the kinematics by a static value of 100ms in Mk-Cs and 70ms in Mk-Jo. These lags were determined by testing the models on various delays to optimize the model performance. We trained and tested the models using the leave-one out cross-validation method: we iteratively set aside one trial for testing and trained the model using the remaining trials. We pulled together kinematic and neural data over different objects (small sphere, large sphere and cylinder) for Mk-Jo and over different positions (left, center and right) for Mk-Cs. The performance was assessed using the coefficient of determination R^2^ for which we computed the 95% confidence intervals across all repetitions. 3D static hand posture reconstruction represents hand configuration for the best time point of the fold resulting in the highest average R^2^ for both the large sphere and the cylinder objects in Mk-Jo.

### Statistics

All computed parameters were quantified and compared between tested groups unless otherwise specified. All data are reported as mean ± SEM unless specified otherwise. Significance was analysed using Kruskal-Wallis test followed by post-hoc multiple comparison analysis between groups.

### Data and software availability

The custom-built open-source software application used to control the robotic arm, along with CAD files of custom objects used with the grip pressure sensor and a step-by-step implementation protocol, have been made available at (<u>10.5281/zenodo.3234138)</u>. Further data from this study are available from the corresponding authors upon reasonable request.

## Author contribution

MC, BB and MB conceived the study; BB designed and implemented the robotic framework hardware and the software modules with the support and supervision of SSM and AB. MB designed and implemented the integrated sensors hardware and software with the support and supervision of FM and SSM. BB, MB, SC, MP and SW conducted the experiments; MB, BB, MP and TM performed the data analysis; SC, MB and MK trained the animals; SC and SW processed the data. MC, MB, BB and MP wrote the manuscript; all authors edited the manuscript; SM and MC supervised the study.

## Acknowledgement

The authors would like to acknowledge the financial support from the Wyss Center grant (WCP 008), the Bertarelli Foundation (Catalyst Fund Grant) an Ambizione Fellowship (No. 167912 to M.C.), and the Swiss National Science Foundation (No. 170032 to SM). The authors would like to thank Prof. Eric Rouiller for his support at the Platform of Translational Neuroscience of the University of Fribourg; Prof. Jocelyne Bloch and Prof. Gregoire Courtine for performing surgical implantation of the BlackRock multi-electrode arrays, Jacques Maillard and Laurent Bossy for the care provided to the animals, Dr. Eric Schmidlin and Simon Borgognon for their help with anaesthesia and surgery preparations, Andre Gaillard for his help with the hardware, and the students Amelie Jeanneret, Laora Marie Jacquemet, Alen Jelusic, Antea Nicora and Clea Kunz for their help processing data.

---

**Figure S1.**
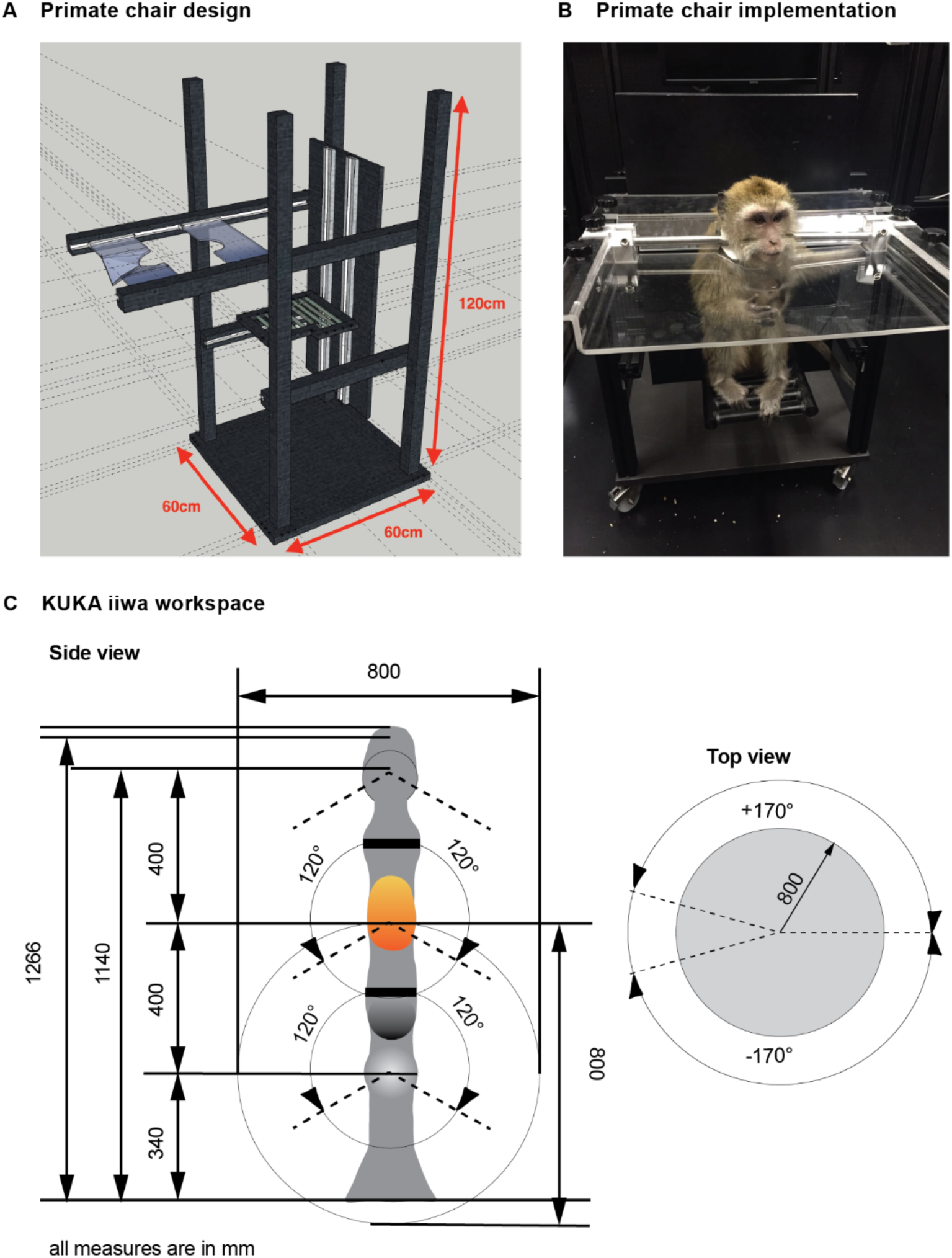
Experimental primate chair and robot workspace. **(A)** Schematic and dimensions of our customized primate chair enabling large ranges of upper limb motion. The height and depth of the seat can be adapted for different animal size. **(B)** Picture of an animal sitting in the chair. **(C)** Dimensions and three-dimensional workspace of the robotic arm.

**Figure S2.**
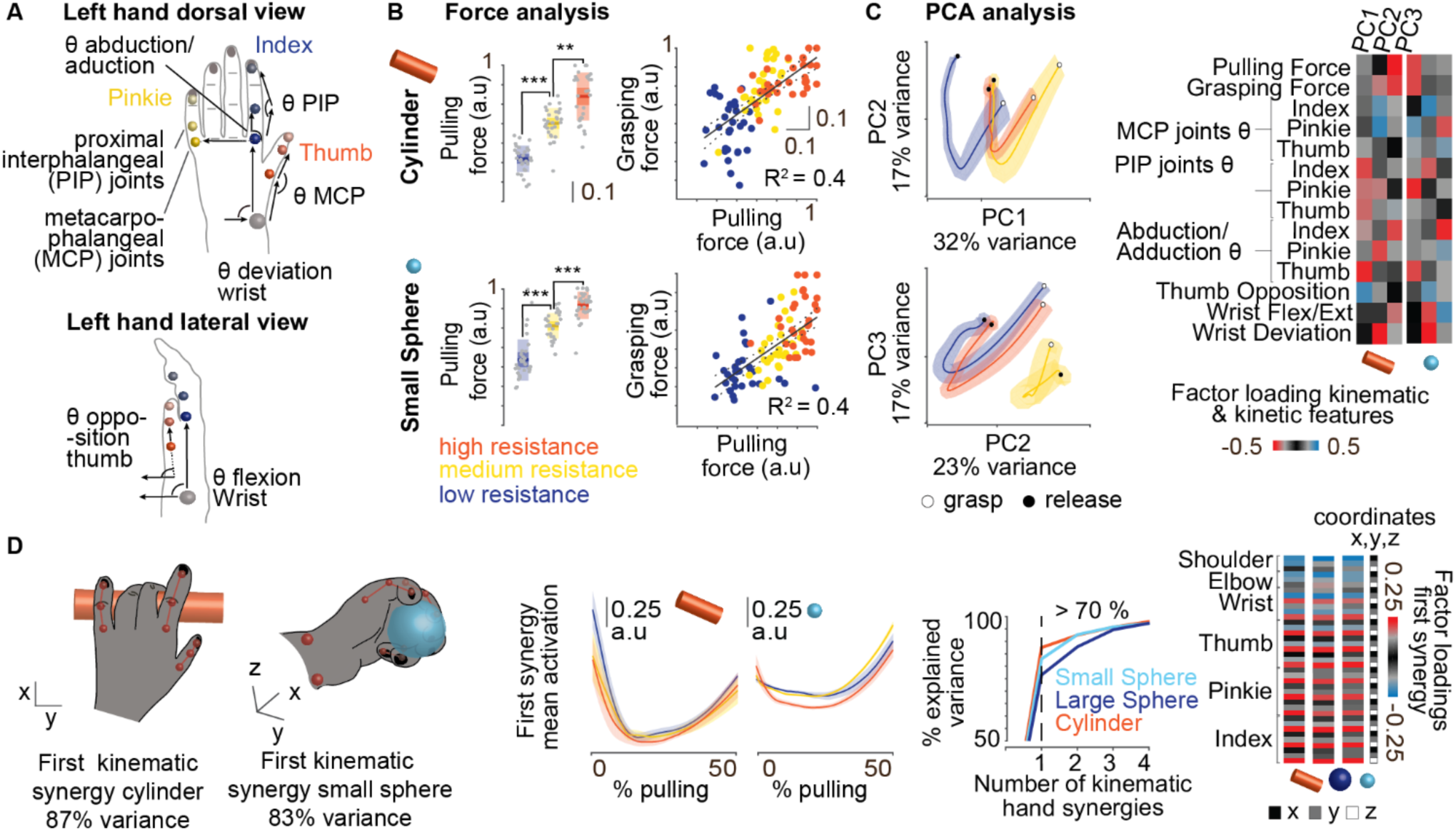
Arm and hand dynamics during natural reaching and grasping. **(A)** Dorsal and lateral view of markers placements and finger joint angles computation for Mk-Jo. **(B)** The box-plots represent the median, the 1.96 SEM and the STD of the pulling force area under the curve (AUC) during small spherical grip and cylindrical grip (n=30 trials per object per condition, three recording sessions). Data were normalized by the maximal pulling force measured within each session. As for the large spherical grip, increase in resistance had a significant effect on the pulling force impulse (Kruskal-Wallis test, 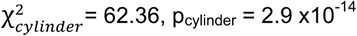, 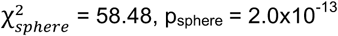 followed by post hoc multiple comparison p_low-med cyl_ = 2.5 x10^-6^, p_low-high cyl_ = 9.6 x10^-10^, p_med-high cyl_ = 0.01 and p_low-med sphere_= 2.0×10^-4^, p_low-high_sphere = 9.6×10^-10^, p_med-high sphere_ = 7.2×10^-4^). Correlation analysis between the grip pressure and the pulling force impulses (AUC) revealed a linear correlation between these two variables (R^2^=0.4 and p_value_ = 5.7×10^-12^ for the small sphere and R^2^=0.4 and p_value_ = 1.2×10^-12^ for the cylinder). **(C)** Principal component analysis of the kinematic and dynamic components of the movement reveals clustering of the three levels of mechanical resistances along the first and second principal components. The color-coded representation of factor loadings identifies variables that contributed most to the differences observed between the difference levels of resistance. **(D)** Postural kinematic synergy defined by the first principal component. The hand postures represented the hand configuration obtained after applying PCA reduction algorithm to the set of hand cartesian coordinates over all trials and representing the loading coefficients of the first PC. Time course activation of the first PC averaged over trials shows modulation for high level of resistance for the small sphere but not for the cylinder suggesting high resistive loads were overcome by changing grip conformation for the small spherical grip but not for the cylindrical grip. Cumulative variance plotted on the right indicates the first kinematic synergy recapitulated more than 70% of the variance for all the grip types. Factor loadings of the first synergies are color-coded on the right showing higher loadings in the x and z directions than in the y direction.

